## Supplementary Figures and Table for "Coordinated multiplexing of information about separate objects in visual cortex"

**Supplementary materials**

Order of Figures/Table:

Figure 1 - Supplementary Figure 1

Figure 1 - Supplementary Figure 2

Figure 1 – Supplementary Figure 3

Figure 2 - Supplementary Figure 1

Figure 3 - Supplementary Figure 1

Figure 4 - Supplementary Table

Figure 7 - Supplementary Figure 1

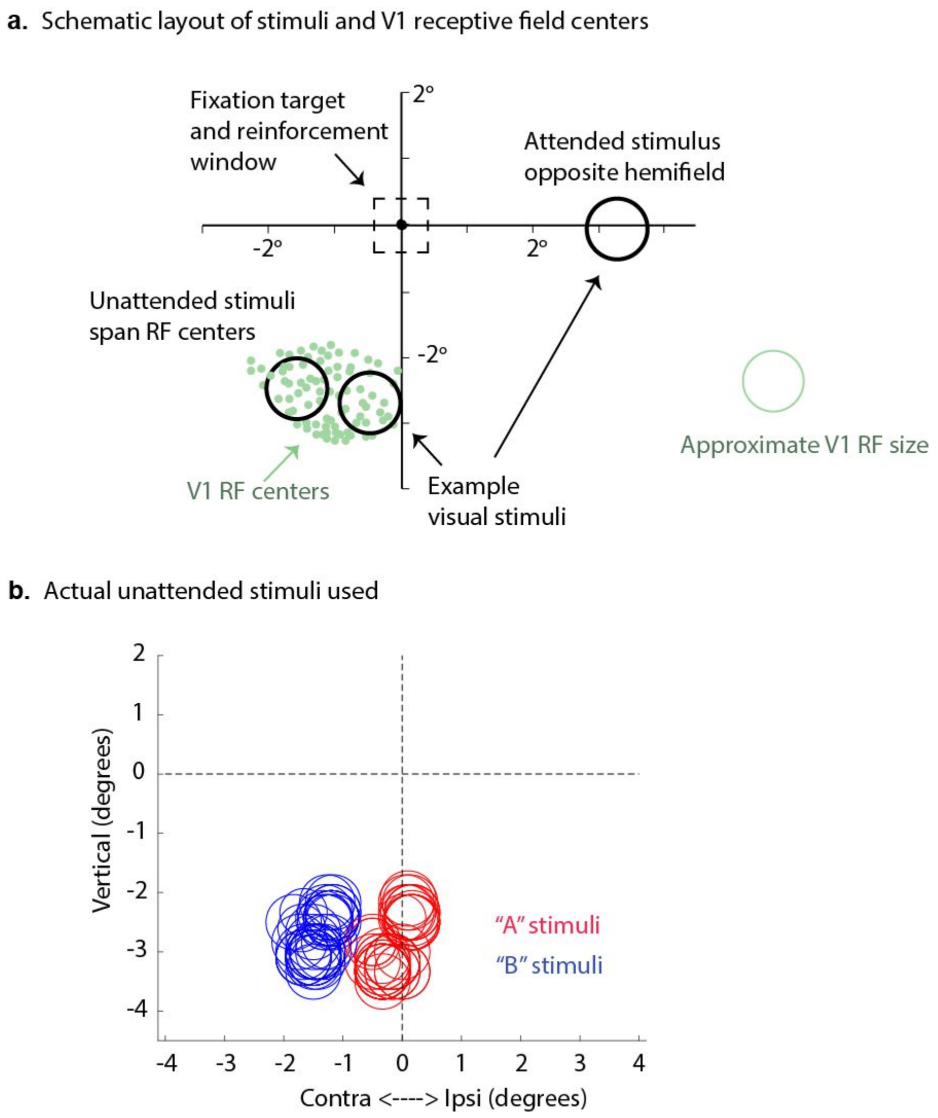

*Figure 1 - Supplementary Figure 1. A. Example receptive field positions, sample estimated receptive field size, and layout of stimuli for the V1 dataset involving adjacent stimuli. This figure is adapted from Figure 1B of Ruff and Cohen 2016. B. Actual stimuli used. One "A" and one "B" stimulus was used in each session. The sizes of the circles indicate the sizes of the Gabor patches; specifically, the radii are equal to twice the standard deviations of the Gaussian envelopes used to construct the patches. Some stimuli were centered slightly into the ipsilateral hemifield but extended into the contralateral hemifield.*

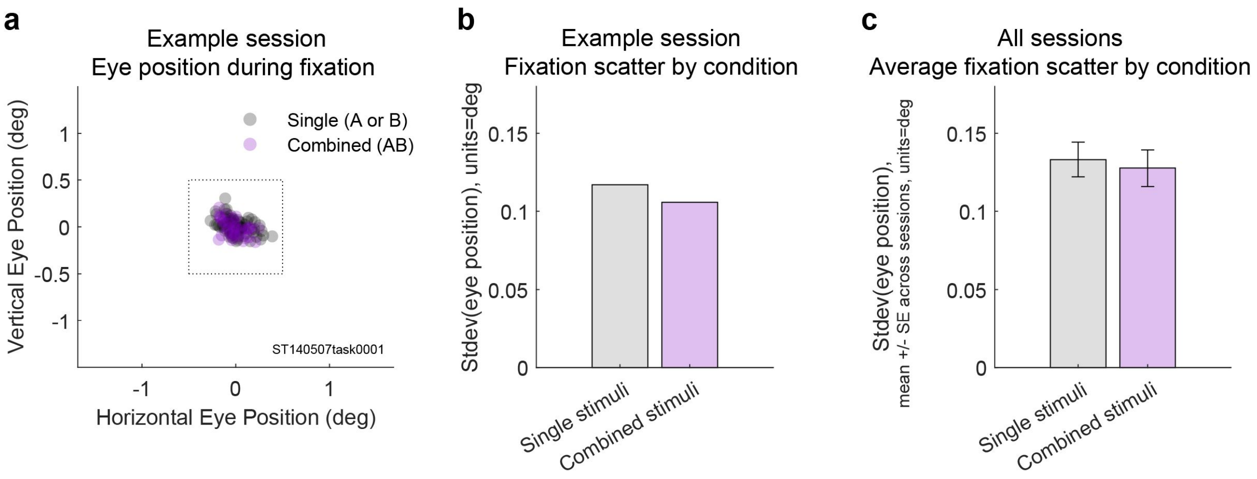

*Figure 1 - Supplementary Figure 2. Eye positions did not differ on single vs. combined stimulus trials (adjacent dataset). A. Average eye positions during stimulus presentations on single (gray) vs. dual (magenta) stimulus trials during one recording session. Box indicates the fixation window, which was +/- 0.5 degrees. B. Geometric mean of the horizontal and vertical standard deviations of eye position for this session. C. Average standard deviation of eye position across all 16 recording sessions as a function of stimulus type. The single vs. combined stimulus values did not differ (one-tailed paired t-test, p=0.927).*

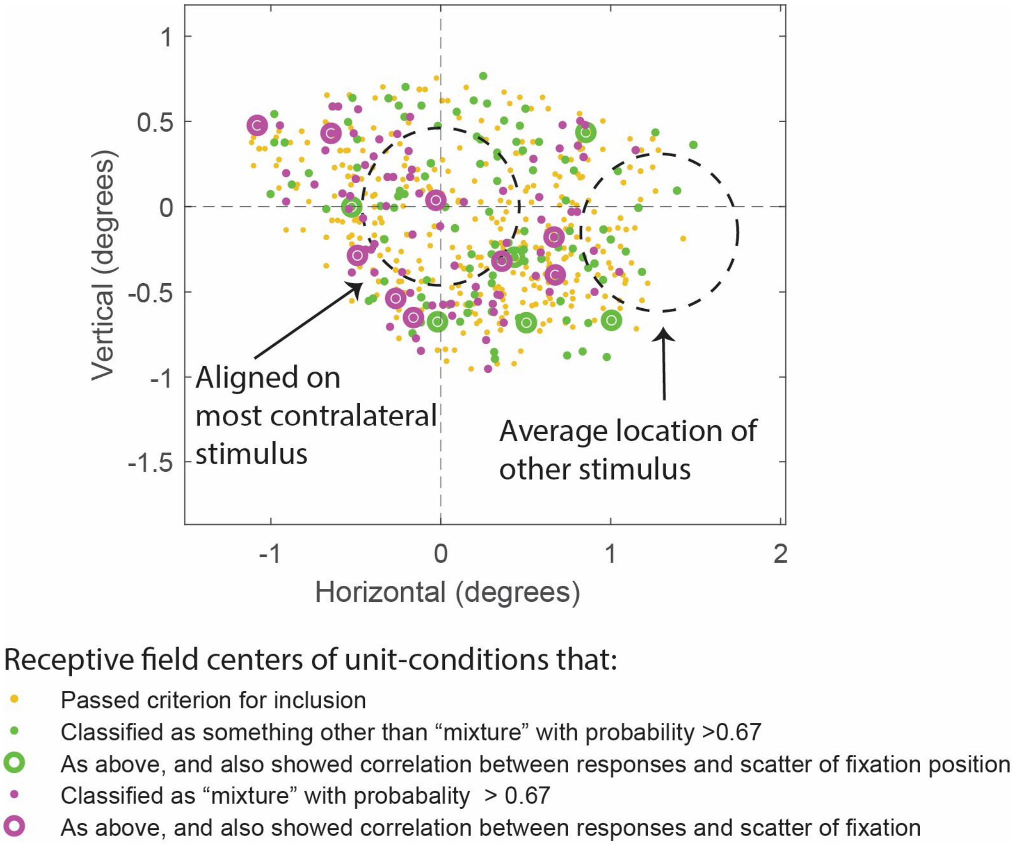

*Figure 1 – Supplementary Figure 3. Relationship between receptive field (RF) location, stimulus location, spike count model classification, and whether firing rates also correlated with scatter in eye position. For clarity, receptive field centers are shown in relationship to the location of the more contralateral of the two stimuli; the other stimulus location is shown as the average of all the other stimulus locations in relation to the most contralateral one. Each dot shows the RF center of one unit-condition (i.e. unit and particular set of stimulus conditions tested in the analysis). Yellow-orange dots correspond to the RF centers for unit-conditions that met criteria for inclusion but for which the Bayesian response pattern classification did not yield a high-confidence result (i.e. probability of successful classification was less than 0.67). Magenta and green symbols indicate RF centers for unit-conditions that were classified as “mixture” (magenta) or as something other than a “mixture” (green) with high confidence (probability of successful classification was greater than 0.67. The magenta and green bull’s eye symbols correspond to unit-conditions that also showed a significant correlation between firing rate and fixational scatter along the dimension connecting the two stimuli for that session (p<0.01). A small number of unit-conditions are not plotted here if the RF mapping did not produce a good estimate of RF center. Overall, significant correlations between spike count and scatter of eye fixation (p<0.01) were identified in 4% of the stimulus conditions that were included for analysis (57/1389, see Table 1). Among the conditions that could be successfully categorized with a probability of at least 0.67 (see Figure 2), the prevalence of significant eye position correlations did not differ significantly for conditions labels as “mixtures” (~9%) vs those that received some other label (~5%). These proportions did not differ from one another by chi-square test (p= 0.1730), nor is there any clear pattern in the location of unit-conditions showing correlations with eye scatter in relation to RF*

*centers and stimulus locations as shown in this figure. Finally, about 9% of “mixtures” (n=9) were responsive to both stimuli (i.e. RF centers were intermediate between the two stimuli and responses exceeded baseline firing by at least one standard deviation for both stimuli alone); among these 9 only 1 showed sensitivity to eye position (11%).*

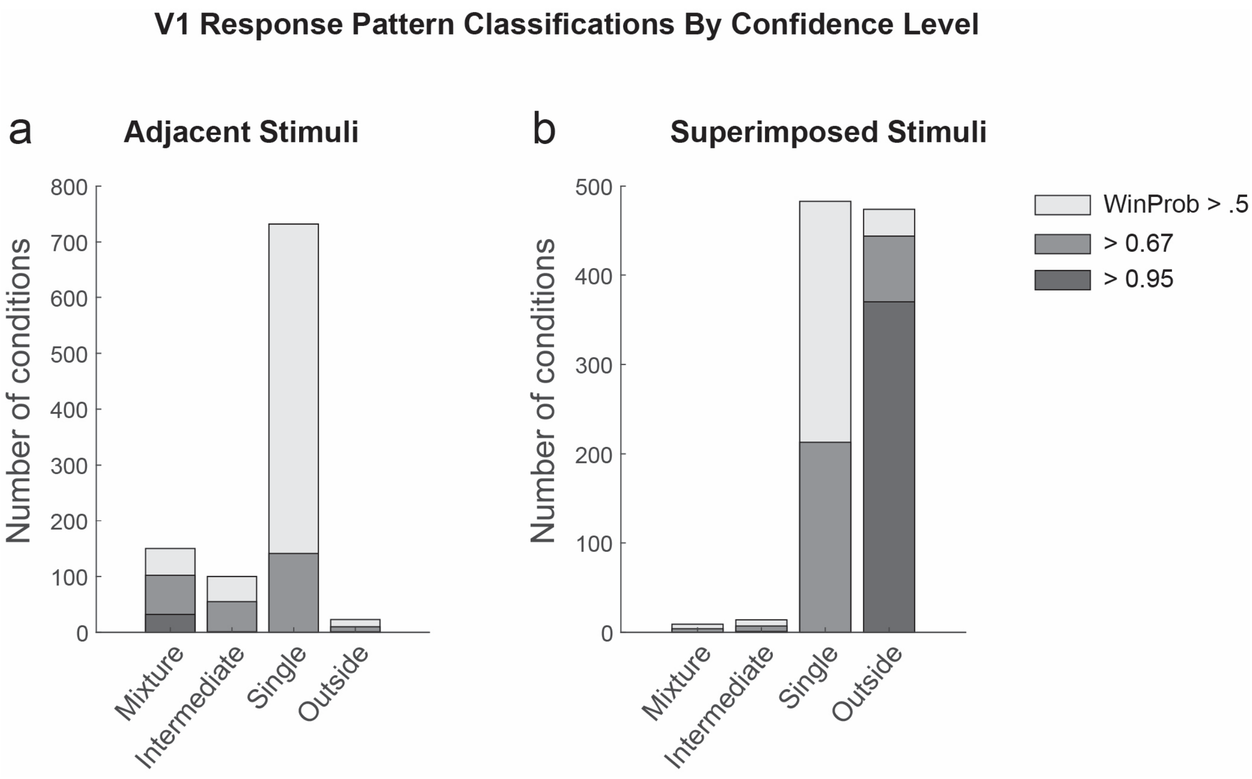

*Figure 2 - Supplementary Figure 1. Detailed results of the response pattern classification analysis on V1 units for the adjacent (A) and superimposed datasets (B). Shading indicates the confidence level of the model categorization. A winning probability of >0.5 indicates that the winning model is at least as likely as all other models combined; a probability of > 0.67 indicates the winner is at least twice as likely as all others, and a probability of 0.95 indicates it is at least twenty times as likely as all others. Cases with a winning probability of >0.25 (the minimum possible) but less than 0.5 are not shown. As noted in the main text, for the adjacent dataset, "mixtures" represented 33% of the cases in which a particular model was at least twice as likely as all other models combined (i.e. the winning probability of 0.67); overall they represented 14% of the cases that passed the exclusion criteria as described in the Methods and Table 1.*

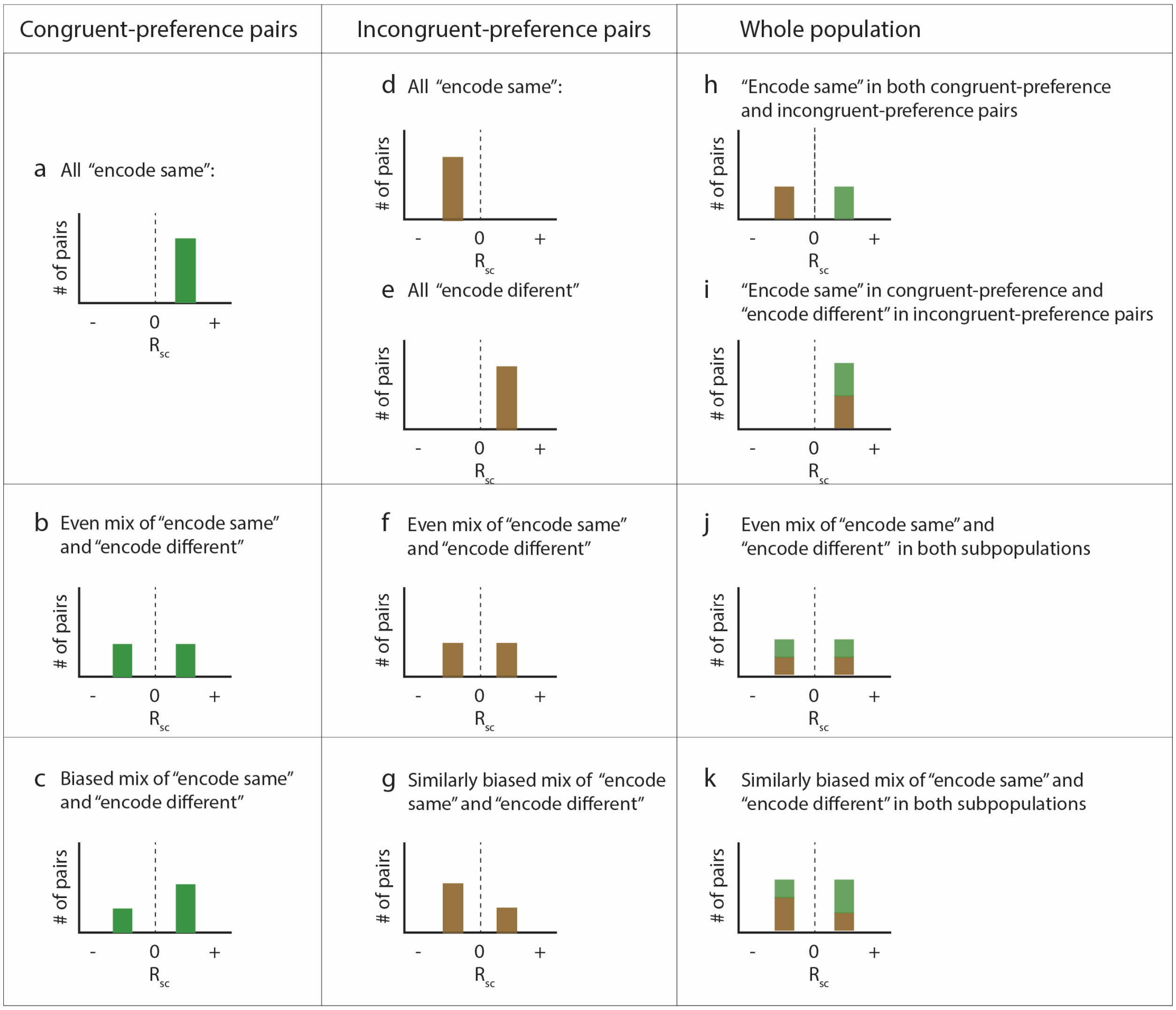

*Figure 3 - Supplementary Figure 1. Details of p*opulation-level predictions under different scenarios. Schematic histograms of the possible patterns of spike count correlations across all the pairs of recorded neurons as a function of whether they share the same *("congruent")* stimulus preferences or have different *("incongruent")* preferences, and whether they tend to "encode" the same stimulus or different stimuli on each presentation (panel h is panel a + panel d; i = a + e; j = b + f; k = c + g; the bottom row (c, g, k) is a biased version of the row above (b, f, j). *Panels h, j, and k are included in Figure 3 as panels d, e, and f.*

| | All | Mixture-Mixture pairs | Intermediate - Intermediate pairs | Single-Single pairs |
|---|---|---|---|---|
| **Congruent AB** | 0.252 | 0.486 | 0.263 | 0.233 |
| **Incongruent AB** | -0.052 | -0.140 | -0.009 | -0.032 |
| **Congruent driven A or B** | 0.251 | 0.384 | 0.266 | 0.250 |
| **Congruent not driven A or B** | 0.145 | 0.132 | 0.142 | 0.147 |
| **Incongruent A or B** | 0.116 | 0.127 | 0.076 | 0.108 |
| **Congruent minus Incongruent AB** | 0.304 | 0.626 | 0.272 | 0.265 |
| **Congruent driven minus incongruent A or B** | 0.135 | 0.257 | 0.190 | 0.142 |

*Figure 4 - Supplementary Table. Median spike count correlations for additional subgroups of the V1 adjacent stimuli dataset. The top two rows show the median spike count correlations observed for dual stimuli for various types of pairs of units, and correspond to the data shown in figures 4 and 5 in the main text (gray background). The next three rows show the same analyses conducted for trials involving single stimuli. Here, the "congruent" group was subdivided according to whether the presented stimulus was the one that elicited the stronger response ("driven") or the weaker one ("not driven"). The bottom two rows show the differences in the medians observed for the relevant congruent and incongruent groups (lines 1 minus 2 and lines 3 minus 5; green background).*

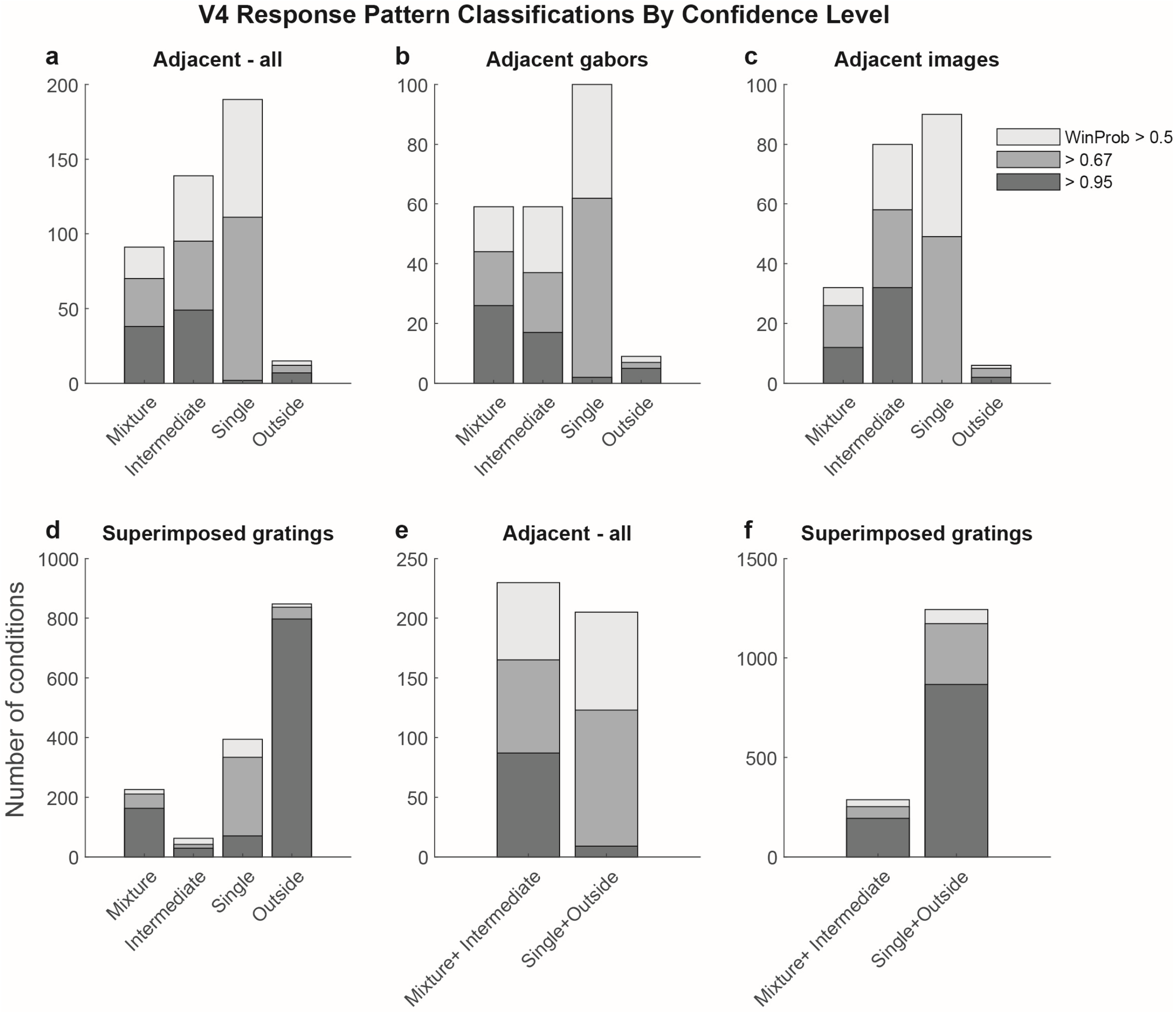

*Figure 7 - Supplementary Figure 1. Detailed results of the whole trial analysis on V4 units for the adjacent (A-C, E) and superimposed datasets (D, F). Shading indicates the confidence level of the model categorization as described in Supplementary Figure 3. Experiments involving adjacent gabors and adjacent images are combined in panels A and E and broken out separately in panels B and C. Panels E and F show the sums of the corresponding bars in panels A and D.*

*Source data file: Source data for the figures and analyses in this manuscript are included as a zip file. The file and folder names are informative regarding which analyses they relate to. Some analyses are based on multiple runs of the modeling code, with slight variations due to the probabilistic nature of the analysis.*
